# MOPRs in mouse islets of Langerhans modulate cell signaling and secretion

**DOI:** 10.64898/2025.12.15.694418

**Authors:** Michael Keith, M Christine Stander, Diego De Gregorio, Andy Huang, Sepideh Sheybani-Deloui, Shannon Townsend, Jeffrey M Zigman, Jing W Hughes, Daniel C Castro

## Abstract

**Article highlights:**

- Mu opioid receptors are expressed on multiple islets of Langerhans cell types
- Mu opioid receptors on islets engage canonical Gαi signaling cascades in islets
- Mu opioid receptors on islets modulate calcium influx and oscillations
- Mu opioid receptors on islets modulate insulin and glucagon secretion.

Most clinically and recreationally used opioids drugs act on the endogenous mu opioid receptor (MOPR). While MOPR is typically studied in the context of addiction and analgesia, decades of evidence indicates that they have a strong modulatory role on metabolism and glycemia. However, whether these effects are directly driven by MOPR actions on pancreatic islets remains poorly understood. Here we sought to comprehensively profile MOPRs on islets to assess how their activity shapes cellular physiology and secretion. First, we used RNA-seq, fluorescent in situ hybridization, and immunoblotting approaches to map islet expression. We observed robust expression of MOPR across multiple cell types in islets. Next, using a FRET-based approach, we show that MOPRs recruit canonical inhibitory pathways, reducing cAMP accumulation. Correspondingly, islets from constitutive MOPR knockout mice showed increased calcium influx and oscillations. However, MOPR knockout had no effect on insulin secretion, instead increase glucagon secretion. Surprisingly, while MOPR antagonism increased overall calcium, it reduced calcium oscillations and suppressed insulin secretion. By contrast MOPR agonism suppressed calcium, increased oscillations, and had no effect on overall hormone secretion. Collectively, these results suggest that MOPR can profoundly shape islet activity, with these effects likely driven by their actions on distinct cell types.

## Introduction

The leading cause of peripheral neuropathy is diabetes, with diabetic neuropathies accounting for approximately 50% of all cases ^1^. While non-opioid analgesics like gabapentin and pregabalin ^2^ are often used as first line treatments for painful neuropathies, opioids such as Tapentadol are frequently used to treat individuals for whom these other drugs are insufficient ^1–4^. Most of these clinically used opioids act through the mu opioid receptor (MOPR). MOPR agonists have been the gold standard analgesic agent for moderate to severe pain for many decades ^5^. The global prevalence of opioid use and diabetes are approximately 50 million and 400 million respectively and are both predicted to increase ^6,7^. As a result, there is a high percentage of individuals that are thus comorbid for both opioid use and diabetes. These individuals have an increased risk of early mortality compared with diabetic non-opioid users ^8^. Further, while the prevalence of type 2 diabetes in individuals abusing opioids is lower compared to non-opioid abusers (partly due to reduced body weight in opioid abusing individuals ^9^) opioid dependent diabetic patients have increased concentrations of hemoglobin A1c compared to nonopioid-dependent diabetic patients, further pointing towards a complex interplay between opioid use and diabetes ^10^.

Decades of evidence indicate that one mechanism through which opioids may modulate diabetes and metabolism is through their direct actions on pancreatic islets. Both exogenous and endogenous opioids have been shown to dysregulate insulin secretion upon systemic administration ^11–18^. Opioid modulation through genetic deletion and pharmacology in-vivo has also been shown to impact glucose tolerance and disrupt feeding behavior in mice ^19,20^. Previous reports ^21^ have also shown that MOPR antagonists increase glucagon secretion from alpha cells in both ex-vivo human and mouse islets of Langerhans. Islet transcription of MOPR is also reduced in humans with type two diabetes and in leptin receptor deficient obese mice (*db/db*) ^21^ indicating that different pathological states are implicated in long term regulation of pancreatic MOPR expression. Given the widespread use of opioids both in medical, illicit and potentially contraindicated contexts, elucidating the cellular and molecular role of pancreatic MOPRs is crucial.

MOPRs are a class A G protein coupled receptor (GPCR) that canonically signals through a Gαi/o mechanism. Upon agonist stimulation, Gαi/o separates from the Gαβγ heterotrimer and inhibits adenylate cyclase to reduce cAMP accumulation while the Gβγ heterodimer binds to G protein inward rectifying K^+^ channels and voltage gated Ca^2+^ channels to increase K^+^ leak and decrease influx of Ca^2+^ ^22^. However, whether these canonical mechanisms hold true within pancreatic islets remains unknown. In this study, we first survey and validate MOPR expression across islet cell types through in-situ hybridization, RNA-seq, and protein expression with immunoblotting. Next, we tested how MOPR constitutive knockout and pharmacological approaches modulated converging second messengers associated with both canonical MOPR signaling and islet hormone secretion such as cAMP and Ca^2+^ accumulation. Finally, we assessed how MOPR impacted insulin and glucagon hormone secretion, thereby creating a direct connection between cellular physiology and functional output. In sum, the data reported here suggests that the MOPR is functionally expressed in islets of Langerhans and can retune cellular physiology and hormone secretion, thus providing a potential mechanism though which opioids could contribute to dysglycemia.

## Methods

### Single cell sequencing

To examine the expression of opioid receptors and peptides in pancreatic islets, we re-analyzed a previously published single-cell RNA-seq dataset of pancreatic islet cells ^23^. This dataset includes 6,523 and 5,924 islet cells from 8-week-old standard chow–fed wild-type (WT) and ghrelin-knockout (GKO) littermate mice, respectively. In this study, we assessed data from WT mice exclusively. After removal of two clusters deemed contaminants from surrounding non-islet tissue, 11 clusters remained, corresponding to the four traditional islet endocrine cell types (α, β, δ, and γ cells, identified by canonical markers) and seven nonendocrine cell populations (endothelial, stellate, and immune cell subsets). Cluster identities and marker genes were previously described 21. Data was processed using Seurat v5. Normalized expression values, generated by the “LogNormalize” method in Seurat, were used for all downstream analyses and visualization. Feature plots were generated to display the distribution and relative expression of selected opioid receptors and peptides across the identified islet cell types.

### Mouse lines

Animals were maintained in accordance with Institutional Animal care and Use Committee (IACUC) regulations at Washington University in St Louis (protocol l#22-0100). Isolated islets from C57BL/6J mice were used in pharmacological experiments and as wild type controls when investigating MOPR knock out islets. Constitutive MOPR knockout mice were originally acquired from Jackson Laboratory and bred in house (strain #007559). All experiments were conducted on 12-week-old mice and sex matched. An equal number of male and female mice were used within independent experiments.

### Fluorescence in situ hybridization

Islet mRNA was imaged using RNAscope™ HiPlex12 assay (ACDbio/Biotechne) and was performed as per manufacturer’s instruction manual. Briefly, mice were transcardially perfused with PBS while anesthetized with isoflurane (to effect). Pancreas were extracted and immersed in 10% neutral-buffered formalin for 16 hours at room temperature. Tissue was removed from formalin and dehydrated through a standard ethanol series. Dehydrated specimens were then cleared and embedded in paraffin. Paraffin blocks were sectioned at 5μm thick and sections placed on Superfrost Plus slides (Thermofisher, 22-037-246). Slides were incubated at 60⁰C for 1 hour, then the paraffin was removed by incubating in 100% xylenes, then 100% ethanol. Slides were air dried, then heat-induced target retrieval was performed to reduce crosslinking and to allow the probes to access the tissue. Slides were dried, again, and treated with protease III for 30 minutes at 40⁰C. Sections were washed and probes for 9 different mRNA targets were added. Probes were allowed to hybridize for 2 hours at 40⁰C. Slides were washed briefly in buffer before performing a series of amplification steps. Once the amplification steps were completed, the fluorophore was added. (Fluorophore ATTO 550 was not used as we saw reduced signal in this wavelength, regardless of expression level.) Slides were counterstained and mounted before being imaged for the first 3 probes.

Slides were incubated in 4XSSC to remove coverslips. Fluorophores bound to the first round of probes were irreversibly cleaved, slides were washed to remove the cleaving agent, then the next round of fluorophores was applied. Slides were counterstained and mounted for imaging the subsequent round of probes. This process was repeated until all probes were imaged. All probes used in this study are included in the table below with the corresponding gene, Catalog # for RNA probe from ACDBio and fluorophore.

| <u>Gene</u> | <u>Probe</u> | <u>Catalog #</u> | <u>Fluorophore</u> |
| --- | --- | --- | --- |
| <u>Oprm1</u> | <u>Mm-Oprm1-T3</u> | <u>315841-T3</u> | <u>ATTO647N</u> |
| <u>Oprl1</u> | <u>Mm-Oprl1-T11</u> | <u>514301-T11</u> | <u>ATTO 647N</u> |
| <u>Oprd1</u> | <u>Mm-Oprd1-T7</u> | <u>427371-T7</u> | <u>ATTO 647N</u> |
| <u>Oprk1</u> | <u>Mm-Oprk1-T8</u> | <u>316111-T8</u> | <u>Alexa Fluor 750</u> |
| <u>Ins1</u> | <u>Mm-Ins1-T9</u> | <u>414661-T9</u> | <u>Alexa Fluor 488</u> |
| <u>Gcg</u> | <u>Mm-Gcg-T4</u> | <u>400601-T4</u> | <u>Alexa Fluor 750</u> |
| <u>Sst</u> | <u>Mm-Sst-T5</u> | <u>404631-T5</u> | <u>Alexa Fluor 488</u> |
| <u>Ppy</u> | <u>Mm-Ppy-T1</u> | <u>437251-T1</u> | <u>Alexa Fluor 488</u> |

### Mouse islet isolation procedure

Adult mice were anesthetized with isoflurane and their pancreases were surgically removed. The mice were sacrificed immediately following pancreas removal via decapitation. The extracted pancreases were washed twice with 8mL Hanks balanced salt solution (HBSS - Gibco, 14025134) and supplemented with 2mg/mL collagenase P (Sigma, 11249002001) for digestion and incubated at room temperature for 35 minutes on tube rotator and inverted 40 times per minute. Following collagenase digestion, the pancreas tissue was washed twice with 10mL HBSS with 1% bovine serum albumin (BSA, Sigma 9048-46-8) at room temperature and strained with a cell strainer. Islets were isolated from strained pancreas tissue by adding 15mL of Histopaque 1100 gradient (Sigma-Aldrich, 10771 and 11191) then centrifuging at 300g for 20 minutes. Following centrifugation, the tissue that remained in the supernatant was washed with HBSS 1% BSA twice and finally resuspended in ‘islet culture media’ consisting of RPMI 1640 media (Thermofisher, 11879020) supplemented with 11mM glucose (Thermofisher, A16828.36) 10% fetal bovine serum (Thermofisher, A52568011) 1% penicillin/streptomycin (Thermofisher, 15140148). After isolation, islets were cultured in 100mm untreated Petri dishes (Thermofisher, FB0875713) overnight at 37^⁰^C, 5% CO_2_ humidified incubator for islet recovery from the isolation procedure. The following day, islets were picked at room temperature under a light microscope for experimental use. Typically, the islet yield per mouse was approximately 150 +/-50.

### Western blot in ex-vivo murine islets

MOPR protein was detected through western blotting using a MOPR specific antibody directed at the C-terminus (7TM antibodies, 7TM0319N) and a GAPDH specific antibody as loading control (Cell Signaling Technologies, 2118). Isolated islets were lysed in 20uL RIPA buffer (Thermofisher, 89900) supplemented with protease cocktail inhibitor (Thermofisher, A32963) and 0.1mM phenylmethylsulfonyl fluoride (PMSF, Sigma Aldrich 10837091001) for 30 minutes on ice, mixing briefly every 10 minutes. Lysates were then separated from cell debris by centrifugation at 10,000g 4⁰C for 15 minutes. Protein content of lysates was determined with Bicinchoninic acid (BCA) protein assay (Thermofisher, 23227). Following determining protein content of the lysate, 20μg of total protein was loaded on to 4%-20% Mini-PROTEAN TGX 10 well gel (BIO-RAD, 4561094) and separated by molecular weight by SDS-PAGE. Following total protein separation, proteins were transferred to PVDF membranes (BIO-RAD, 1620264) at 100V for 2 hours at 4⁰C. Membranes were then blocked with ‘EveryBlot’ blocking buffer (BIO-RAD, 1620264) for 30 minutes at room temperature under constant agitation. Membranes were incubated at 4⁰C overnight under constant agitation with primary antibodies, either anti-MOPR (7TM antibodies, 7TM0319N) or anti-GAPDH (Cell Signaling Technologies, 2118S) for loading control. The following day, the membranes were washed four times with TBS-T for 10 minutes each and incubated with secondary anti-rabbit Horseradish peroxidase antibody conjugate dissolved in TBS-T and supplemented with 2% fat-free milk powder. The membrane was subsequently washed 5 times for 10 minutes each with TBS-T, 10 to remove unbound secondary antibody and visualized by mixing enhanced chemiluminescent (ECL) substrate 1:1 as per manufacturer directions (BioRad, 1705060). After a two minute ECL incubation, the membrane was flicked dry and chemiluminescence was imaged using a ChemiDoc imager (BioRad, 12003153).

### cAMP FRET

Plate-based cAMP measurements were made using the LANCE Ultra cAMP kit (Revvity, TRF0262). Islets were picked into a 1.5mL Eppendorf tube and washed twice with HBSS 1% BSA then resuspended at 1 islet/μL in stimulation buffer consisting of HBSS, 1% BSA, 0.5mM IBMX (Thermofisher, PHZ1124) and 5mM HEPES (Thermofisher, 15630106). 5uL of 1 islet/μL stimulation buffer was dispensed into each assayed well of a 384 low volume proxi-plate (Revvity, 6008280) and incubated with additional stimulation buffer supplemented with appropriate glucose concentration for the specific experimental parameters, then covered with plastic plate covers and equilibrated to the desired glucose concentration through incubation in a 37⁰C, 5% CO2 humidified incubator for 45 minutes. If applicable to the experiment, following glucose equilibration, islets were stimulated with forskolin or vehicle control at the indicated concentration of the specific experiment for 20 minutes at room temperature. After forskolin stimulation, islets were treated with 10μM selective MOPR antagonist CTAP (Sigma-Aldrich, C6352) or equivalent vehicle for 10 minutes at room temperature. Following antagonist treatment, islets were treated with 10μM selective MOPR agonist DAMGO (AChemBlock, 78123-71-4) or equivalent vehicle for a further 10 minutes before initiating the end point of the assay by adding the detection reagents; Europium cryptate-cAMP and uLight-anti-cAMP conjugated antibody which are both diluted in cell lysis buffer supplied by the manufacturer as described in the assay kit. Following detection reagent addition, the plates were incubated for 1 hour at room temperature, in the dark. In tandem to islet trials, an exogenous cAMP standard curve was also established on the same plate as the LANCE assay kit instructions indicated. Following the final hour of detection reagent addition, homogenous time resolved fluorescence (HTRF) was read at 665nm/615nm with 320nm excitation on a Flexstation 3 plate reader (Molecular devices). Each data point reflects the mean from 4 technical replicates per mouse.

Ca^2+^ imaging.

All live-cell Ca^2+^ confocal imaging was conducted using a Zeiss Airy scan LSM 880 confocal microscope (Zeiss). to visualize Ca^2+^, 20-30 islets were handpicked under a light microscope into an untreated 35mm Petri dish (CellTreat, 229638) and incubated in islet culture media with 4μM Calbryte 520 AM cell permeable Ca^2+^ dye (AAT Bioquest, 20653) for 75 minutes. Following Calbryte incubation, the islets were picked again and mounted on a human recombinant laminin-521 (Thermofisher, A29248) treated 35mm glass imaging dish (Mattek corporation, NC9341562) filled with 2mL Krebs-Ringer bicarbonate HEPES (KRBH) buffer consisting of (128.8 mmol/L NaCl, 4.8 mmol/L KCl, 1.2 mmol/L KH2PO4, 1.2 mmol/L MgSO4⋅7H2O, 2.5 mmol/L CaCl2, 20 mmol/L HEPES), 5 mmol/L NaHCO3, and 0.1% BSA [pH 7.4]) at the desired glucose concentration depending on the experiment for 90 minutes. Following islet adherence, the glass dishes were imaged within a chamber maintained at 37⁰C, 5% CO2 and allowed to reach equilibrium in the chamber for 10 minutes. After the equilibration, islets were located using brightfield and subsequently imaged live using a 488nm Argon laser for excitation and emission was detected at 500-570nm using a spectral detector. All images were captured using Plan-Apochromat at 10x magnification/numerical aperture 0.45 with a frame size of 512×512 (135μm^2^ x 135μm^2^), a pinhole size of 206.6μm, a pixel dwell time of 2.05 μ seconds and a scan time of 2.52 seconds. All images were captured at 820 master gain and 0.40 laser power across all imaging experiments. Continuous images were taken every 2.52 seconds for a minimum of 300 total image cycles at 3mM glucose to establish a baseline then a minimum of 800 total image cycles per experimental condition conducted at 9mM glucose. For experiments in which glucose concentration was increased from 3mM to 9mM, KRBH supplemented with glucose was added by manual pipetting and quickly reimaged to capture the low to high glucose transition. In Ca^2+^ imaging experiments assessing 3mM to 9mM glucose concentration transitions, drug controls (CTAP or DAMGO) were added to the imaging plate concurrently with glucose. When assessing drug effects on Ca^2+^ oscillations, islets were equilibrated at 9mM glucose KRBH for one hour and imaged for 1000 cycles before adding 10μM DAMGO or CTAP. Following drug addition, islets were imaged for a further 1000 cycles.

Images were analyzed using FIJI/ImageJ and region of interest (ROI) analysis was conducted on individual islets to capture whole islet signal intensity across the entire time series. In experiments investigating the 3mM to 9mM glucose transition, ROI signal was normalized to the 3mM glucose baseline through subtraction and each islet from an individual mouse and independent experiment were averaged where every data point is the mean of islets originating from an individual mouse. In oscillation analysis experiments requiring single islet resolution for interpretation, each data point reflects individual islets from at least n=6 age matched mice (3 female, 3 male).

### Static secretion assay

For constitutive MOPR KO versus wild type control experiments, secretion samples were taken sequentially at different glucose concentrations from the supernatant of adherent isolated islets. Initially, islets were equilibrated to KRBH buffer supplemented with 2.8mM glucose for 45 minutes at 37°C and 5% CO_2_ and an initial baseline secretion sample was collected by aspirating the supernatant. The 2.8mM glucose KRBH buffer was then replaced with 3mM glucose KRBH and islets were incubated for an additional 45 minutes at 37°C and 5% CO_2_ to obtain low glucose condition samples. The glucose concentration was then increased to 9mM by aspiration and replacement with 9mM glucose KRBH and the islets were incubated for another 45 minutes at 37°C and 5% CO_2_ before the high glucose condition sample was taken.

For secretion assays investigating pharmacological modulation of MOPR, islet secretion samples were taken non-sequentially. Distinct groups of islets from the same mice were incubated separately and at 3mM and 9mM glucose. MOPR antagonist CTAP and MOPR full agonist DAMGO were added at the start of the 45-minute incubation with the final glucose concentration where either 3mM or 9mM glucose was assessed. Each drug was tested at 1µM and 10µM concentration. To extract total islet hormone content, islets were incubated in acid-ethanol (1.5% HCl, 70% ethanol) for at least 24 hours at −20⁰C. All samples were stored at −20⁰C until ELISA was run to quantify hormone secretion.

### Insulin ELISA

Islet samples were thawed from −20⁰C on ice and the ELISA was conducted as instructed in the Crystal chem Ultra-Sensitive Mouse Insulin ELISA kit (CrystalChem, 62100). Insulin secretion and total insulin content was normalized by using the wide range insulin standard provided with the assay kit. Samples and insulin standards were read on a BioTek synergy neo2 plate reader (Agilent) with Gen5 software measuring absorbance at 450nm and 630nm. The data was then normalized by subtracting the absorbance at 630nm from absorbance at 450nm and insulin concentrations were interpolated from the standard curve of known insulin concentrations.

### Glucagon Lumit

Glucagon secretion was measured using the NanoLuc luminescence based Lumit^TM^ Glucagon immunoassay (Promega, W8022) and performed as described by the manufacturer. In short, secretion samples were thawed as before and dispensed into a 96 well plate and both the Lumit^TM^ anti-glucagon antibody large fragment and the Lumit^TM^ small anti-glucagon fragment were added to every well and incubated for 1 hour at room temperature in the dark. After glucagon antibody fragment incubation, the detection substrate included in the kit was added to every sample and incubated for 5 minutes and luminescence was read on a BioTek synergy neo2 plate reader (Agilent) with Gen5 software. Alongside secretion samples, a glucagon positive control included in the kit was also assessed in every experiment.

## Results

### The Mu opioid receptor is expressed in murine islets of Langerhans

RNA expression of the *Oprm1* gene has been previously validated in both human and mouse islets ^19,21^. Here, we sought to more fully characterize the opioid system in endocrine pancreas using a repertoire of complementary approaches. First, we reanalyzed a previously published single-cell RNA-seq dataset of mouse pancreatic islets to assess opioid signaling components. We found that *Oprm1* was expressed across all major endocrine cell types (Figure 1A). Among opioid peptides, *Penk* and *Pomc* were also detected in subpopulations of α, β, δ, and γ cells. (Figure 1A). We found that *Oprm1* was detected in every pancreatic endocrine cell-type in both male and female islets, as well as evidence for delta and nociceptin opioid receptor expression. Likewise, preproenkephalin (*Penk*), propopiomelanocortin (*Pomc*), preprodynorphin, and prepronocicpetin were also detected in subpopulations of all islet cell types. To further validate *Oprm1* transcription, we used an iterative in-situ hybridization approach to map the co-localization of *Oprm1* with known islet cell type specific markers (Figure 1B). In addition to detecting *Oprm1* transcripts we also found evidence of *Oprl1* and *Oprd1* (Figure S1). Quantification with HALO software revealed that *Oprm1* transcripts were expressed in approximately half of cells within the islets imaged (53%). This expression was proportionally stratified across alpha (Gcg), beta (Ins1), delta (Sst), and gamma (Ppy) cells, such that half of each cell type co-expressed *Oprm1* transcripts. Figure 1C displays the absolute percentage of each cell type, as well as the absolute co-expression with *Oprm1*; 42% of cells expressed Ins1 and *Oprm1*, 5% of cells expressed Gcg and *Oprm1*, 4% of cells expressed Sst and *Oprm1* and 5% of cells expressed Ppy and *Oprm1*. Despite having strong evidence for MOPR expression with RNA approaches, protein expression of MOPR has never to our knowledge been validated in the islets. To validate protein expression of MOPR in the pancreatic islet, we used immunoblotting to detect MOPR protein using a C-tail directed antibody for the MOPR in ex-vivo murine islet lysates. Prior to ex-vivo islet trials, we utilized a HEK293 cell line transiently transfected with either the MOPR or equivalent empty vector control to validate antibody specificity. In these experiments we only observed band development in cells transfected with the MOPR (Figure S2). Following the determination of MOPR antibody specificity in an over-expression system, we sought to probe MOPR protein expression in ex vivo murine islets. In wild-type islets, we detected bands of the same molecular weight as monomers of MOPR, likely at two different glycosylation states. We also detected bands at double the molecular weight of where we suspect the monomer to be, indicating dimeric MOPR. To fully validate MOPR protein expression and confirm constitutive knockout of MOPR for future experiments, we also replicated this immunoblotting experiment in islets isolated from constitutive MOPR KO mice and saw no MOPR protein band development with equivalent protein loaded and identical chemiluminescence development visualization parameters (Figure 1C).

**Figure 1.**
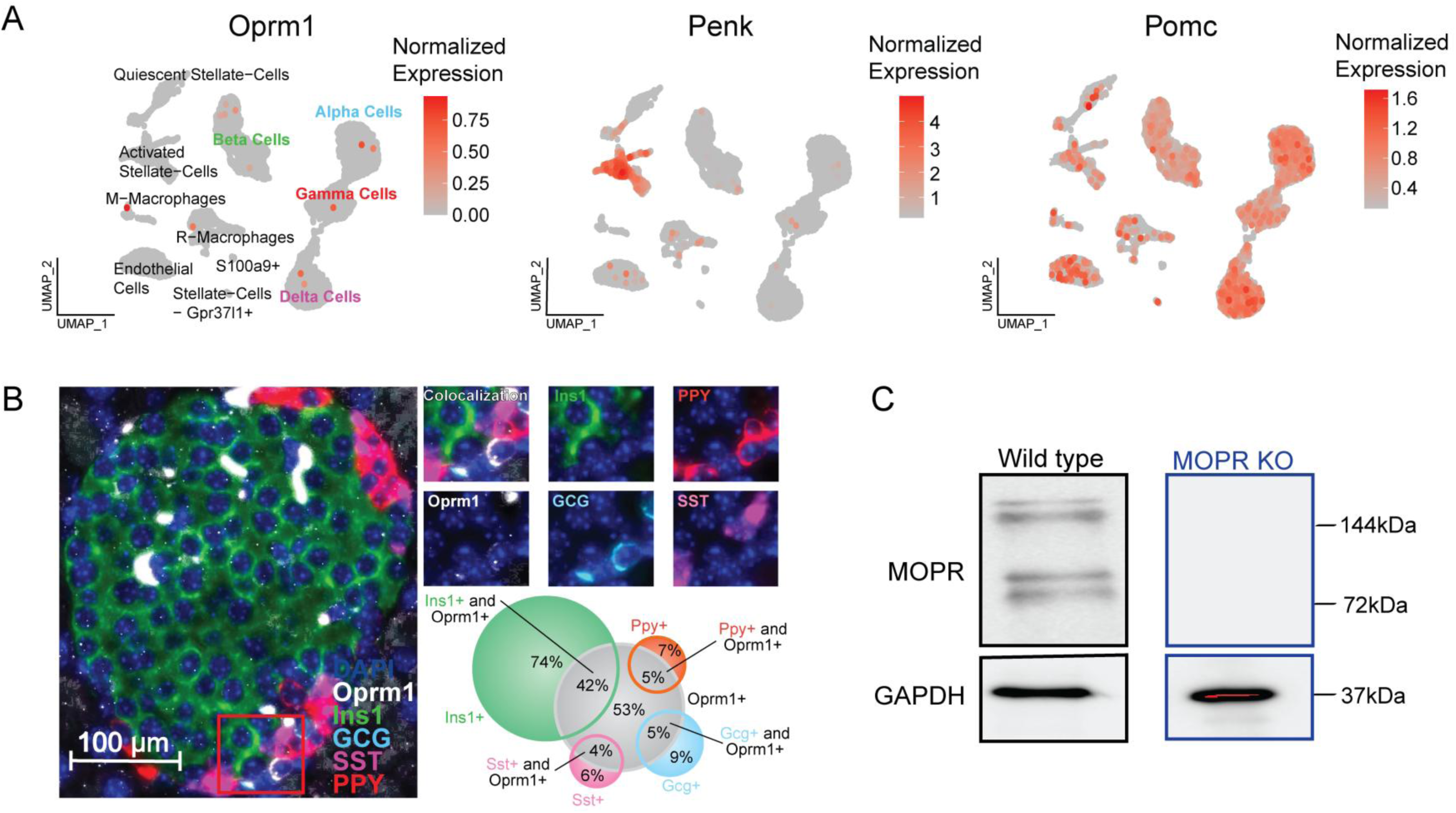
Mu opioid receptor is expressed in mouse islets of Langerhans. A) Feature plots of Oprm1, Penk, and Pomc expression in islet cells (scRNA-seq, GSE244390). Each point is a single cell; color scale shows normalized expression (log-normalized counts). Oprm1, Penk and Pomc are transcribed across all major endocrine cell types. B) Isolated wild type islet RNA scope image and HALO quantification of probe co-localization showing individual and merged transcripts of the mu opioid receptor, (Oprm1, white) insulin, (Ins1, green) glucagon, (Gcg, blue) somatostatin, (Sst, pink) and polypeptide Y (PPY, red) Scale = 100μm MOPR mRNA is expressed in approximately half the islet populations assessed and is co-expressed in all islet cell type specific probes. MOPR mRNA (Oprm1) was expressed in 53% of islet cells, 42% of cells were positive for both Oprm1 and Ins1, 5% of cells were positive for both Gcg and Oprm1, 4% of cells were positive for both Oprm1 and Sst, and 5% of cells were positive for both Ppy and Oprm1. C) Representative western blot images of either wild type ex-vivo isolated murine islets (left panels) or MOPR KO ex-vivo isolated murine islets (right panels) immunoblotted for the Mu opioid receptor (upper panel) and GAPDH for loading control (lower panel). Blots were loaded with 20μg total protein, determined with BCA protein assay.

### Mu opioid receptors are functional in the mouse islet of Langerhans and impact glucose induced cAMP suppression dynamics

After validating expression of MOPR in the islet of Langerhans, we sought to characterize their pharmacological responsiveness by utilizing a cAMP homogenous time resolved fluorescence FRET assay, a schematic of this assay is shown in figure 2A. Typically, the use of cAMP accumulation assays to assess G protein coupled receptors which suppress adenylate cyclase activity (such as Gαi/o coupled receptors like MOPR) require a baseline stimulation of adenylate cyclase to observe suppressive effects of agonists. In this vein, we established a forskolin concentration response curve in ex vivo islets. We observed a typical sigmoidal concentration response curve where higher concentrations of forskolin drastically reduced signal (indicating increased cAMP accumulation). As the concentration was reduced, the magnitude of islet cAMP generation decreased until it reached a plateau where forskolin was no longer detectably impacting cAMP concentration (Figure 2C). To investigate how MOPRs on islets influence islet cAMP, we tested ex vivo islets in 4 conditions: vehicle control, 100nM forskolin alone control, 100nM forskolin with 10μM DAMGO (a selective MOPR full agonist), and 100nM forskolin stimulation with 10μM DAMGO and 10μM CTAP (a selective MOPR antagonist). As expected, forskolin decreased signal compared with vehicle control. In contrast, the addition of DAMGO significantly suppressed forskolin induced cAMP accumulation resulting in approximately 2-fold increase in HTRF 665nm signal. CTAP was able to block the suppressive effect of DAMGO resulting in no significant difference compared to forskolin alone (Figure 2D). Since this particular FRET-based cAMP accumulation assay has not to our knowledge been used in ex-vivo islets, we also replicated this pharmacology experiment in an expression system using HEK293 cells transiently transfected with MOPR. In this experiment we similarly observed decreased signal when cells were treated with forskolin compared with vehicle control as well as the suppressive effect of DAMGO on cAMP accumulation. Analogously to previous cAMP accumulation assays using ex-vivo murine islets, HEK293-MOPR cells treated with 10μM CTAP, 10μM DAMGO and forskolin were not significantly different to treatment with forskolin alone (Figure S3). The results from figure 2D suggest that ex vivo islet cAMP accumulation can be modulated with MOPR pharmacology, indicating functional expression of MOPR and mobilization of associated Gαi/o signaling cascades.

**Figure 2.**
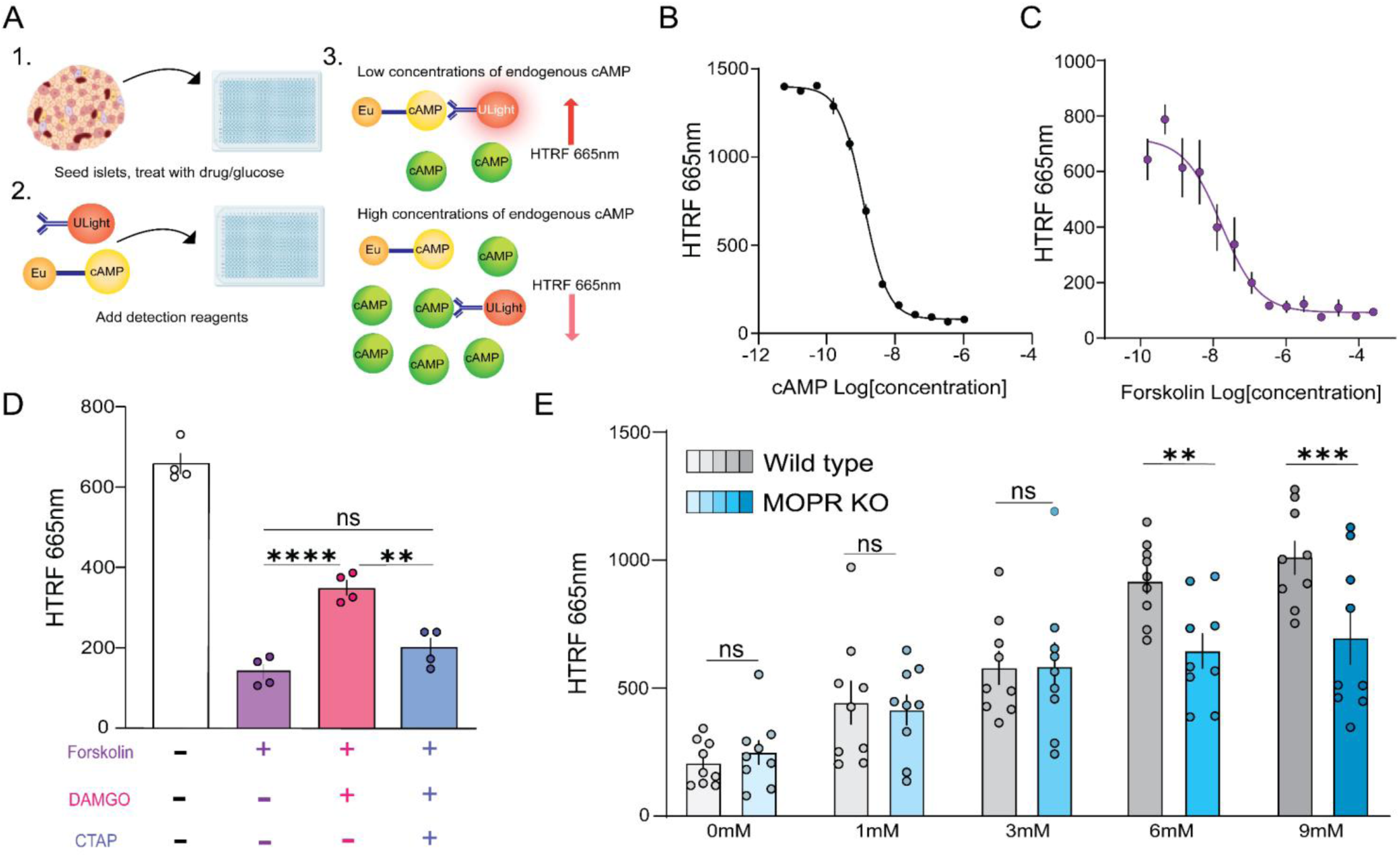
Mu opioid receptors are functional in the mouse islet of Langerhans and mu opioid receptor knockout increases cAMP accumulation under high glucose conditions. A) Assay schematic, 5 islets are seeded to a 384 well plate, treated with appropriate drug or glucose additions contingent on the experimental condition. Then incubated with cAMP detection reagents. Homogenous time resolved fluorescence signal at 665nm is inversely proportional to cAMP concentration B) Representative 12-point cAMP standard curve with exogenous cAMP concentrations starting at 1μM serially diluted 3-fold, n=4 error bars showing +/-SEM C) 14-point Forskolin concentration response curve on ex-vivo isolated islets (5 per well) starting concentration 250μM serially diluted 3-fold, n=5 mice with error bars showing +/- SEM D) Ex vivo murine islets (5 islets per well) subjected to either vehicle control (white), 100nM forskolin (purple), 100nM forskolin and 10μM DAMGO (pink) or 100nM forskolin, 10μM DAMGO and 10μM CTAP (blue). DAMGO significantly suppresses Forskolin’s stimulatory effect on cAMP accumulation; Forskolin vs forskolin + DAMGO ****P<0.0001. CTAP can block DAMGOs action on cAMP accumulation suppression, Forskolin + DAMGO vs Forskolin + DAMGO +CTAP **P<0.01. Due to CTAP blocking DAMGOs effect, there is no significant difference between Forskolin alone and Forskolin, DAMGO and CTAP together, Forskolin vs Forskolin + DAMGO + CTAP not significantly different P>0.05 assessed by one way ANOVA with Bonferroni’s test for multiple comparisons, each data point shows the mean from one mouse n=4, with error bars showing +/-SEM. E) Wild type (greyscale) or MOPR KO islets, (blue scale) 5 islets per well, subjected to a titration of glucose for 45 minutes. MOPR KO has no effect at 0, 1 or 3mM glucose n=9 mice P>0.05 but significantly decreases HTRF 665nm signal at 6mM **P<0.01 and 9mM glucose ***P<0.001, assessed by two-way ANOVA with factors of genotype and glucose concentration. Bonferroni’s test was used to correct for multiple comparisons. Each data point is the mean from islets isolated from n=9 mice with error bars showing +/-SEM

Previous studies have indicated that one of the modalities islets detect physiological changes in glucose concentrations is through modulating cAMP accumulation to tune hormone secretion ^24–27^. In this vein we sought to explore the relationship between glucose concentration and islet intracellular cAMP accumulation. To investigate this, we subjected isolated islets to a titration of glucose concentrations and measured consequent changes in cAMP activity. We found that cAMP accumulation was inversely proportional to glucose concentration, where the highest endogenous cAMP accumulation occurred at lower concentrations of glucose. These levels then decreased as the glucose concentration was increased. When we replicated the experiment using MOPR constitutive knockout islets, this relationship was significantly disrupted. Specifically, HTRF 665nm signal derived from MOPR KO islets were significantly decreased at both 6mM and 9mM glucose concentrations relative to wild type islets (Figure 1E). This data indicates that MOPR biology has some physiological control over the critical relationship between islet glucose responsiveness and subsequent glucose induced intracellular signaling.

### Constitutive knockout of the Mu opioid receptor increases Ca2+ accumulation at low glucose, when transitioning from low to high glucose states and Ca2+ oscillation frequency

Our results thus far have indicated that MOPR is expressed across the islet and is pharmacologically and potentially biologically significant. In particular, the effects of MOPR appear to be glucose concentration dependent, potentially suggesting that it may influence other glucose-dependent islet activity such as Ca^2+^ dynamics ^27–30^. Since MOPR activation canonically leverages Gβγ signaling to increase K^+^ leak through G protein inward rectifying channels and reduce influx of Ca^2+^ through inhibiting voltage gated Ca^2+^ channels, ^22,31,32^ there is potential for a functional intersection between MOPR signaling and glucose dependent islet activity. To investigate this, we implemented a live-cell confocal microscopy approach using Calbryte^TM^ 520 AM Ca^2+^ sensitive dye ^33^ (Figure 3A). It is well substantiated that increased glucose concentrations are positively correlated with intracellular Ca^2+^ accumulation in islets^34–36^, which we also observed (Figure 3B). With this approach, we continuously monitor intracellular Ca^2+^ dynamics in the islet to probe how modulation of MOPR pharmacologically or through genetic deletion impacts islet Ca^2+^ dynamics. First, we found that MOPR KO islets had increased basal Ca^2+^ signal at 3mM glucose compared to wild type islets (Figure 3C). When islets were transitioned from 3mM to 9mM glucose, we found that both wild type and KO islets had increased accumulated Ca^2+^. However, this effect was more dramatic in KO islets, even when controlling for their relatively high baseline (Figure 3D). After the transition to 9mM glucose, KO islets continued to have greater Ca^2+^ accumulation which persisted for the duration of the experiment and resulted in a significantly increased area under the curve (Figure 3E-F).

**Figure 3.**
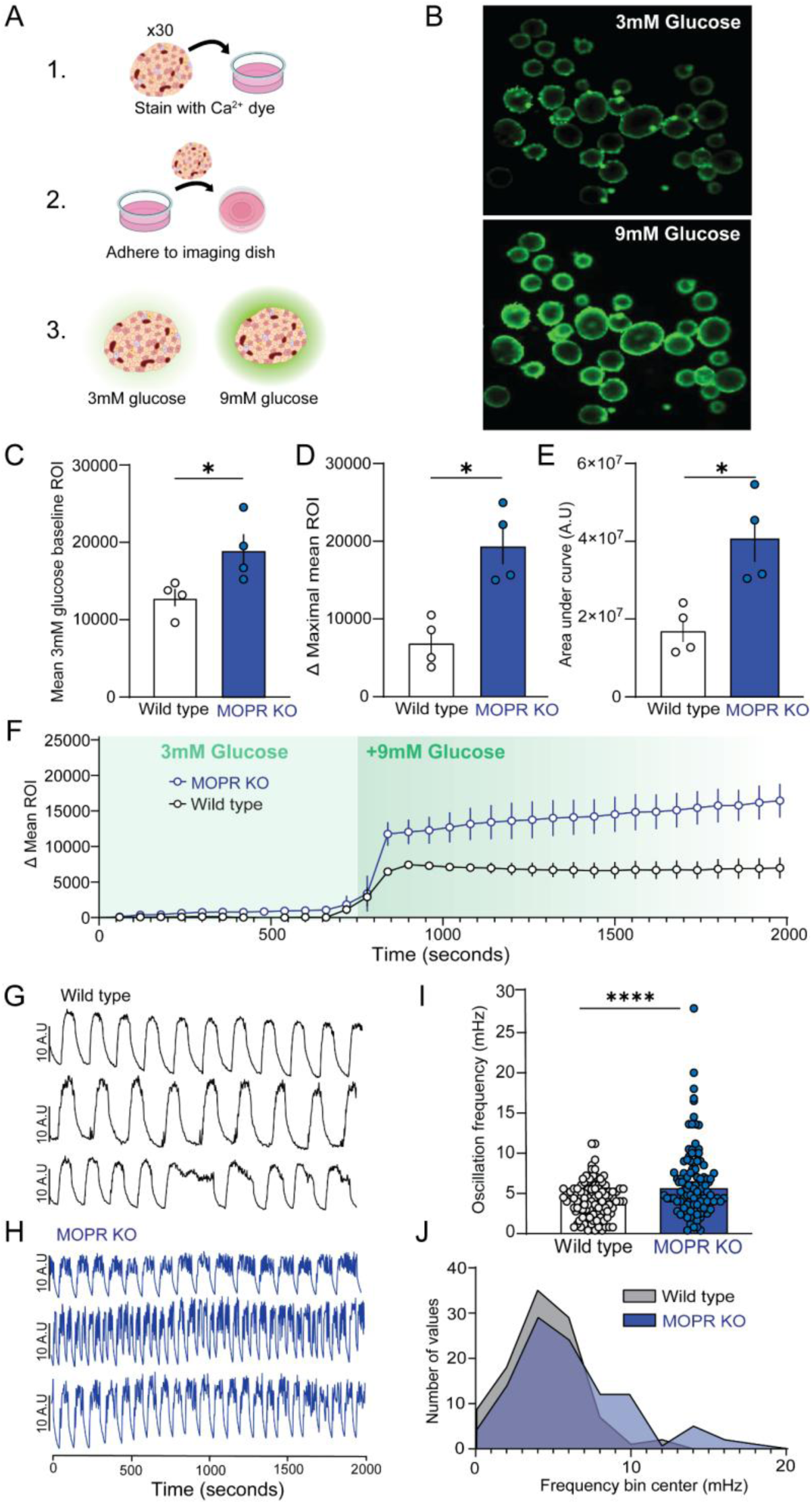
Constitutive knockout of the mu opioid receptor increases Ca2+ accumulation and Ca2+ oscillation frequency. A) Assay schematic and workflow. Islets are stained with Ca2+ dye Calbryte 520AM and adhered to 35mm glass bottom imaging dishes with recombinant laminin. Under the low glucose conditions of 3mM, fluorescence is lower relative to the high glucose condition of 9mM. B) Representative live confocal images of wild type islets at 3mM and 9mM glucose. C) Baseline raw ROI signal at 3mM glucose of both wild type and MOPR KO islets. MOPR KO significantly increased baseline Ca2+ accumulation at 3mM glucose n=4 mice of each genotype, assessed by unpaired t-test *P<0.05, error bars show +/- SEM. D) Maximal delta signal of Ca2+ accumulation through increasing glucose concentration from 3mM to 9mM of both wild type and MOPR KO islets derived from F. MOPR KO significantly increased Ca2+ accumulation, assessed by unpaired t-test *P<0.05 from 20-25 islets per mouse n=4 mice of each genotype, error bars show +/- SEM. E) Area under curve analysis of wild type islets compared with MOPR KO islets, the area under the MOPR KO curve was significantly increased compared with wild type, n=4 mice of each genotype, assessed by unpaired t-test *P<0.05, error bars show +/- SEM. F) Ca2+ accumulation time course starting at 3mM glucose transitioned to 9mM at 750 seconds comparing wild type and constitutive MOPR KO ex vivo islets through mean region of interest (ROI) analysis of 20-25 islets per mouse from n=4 mice of each genotype with delta ROI normalized to signal from 3mM glucose at equilibrium. Error bars show +/- SEM. G) Three representative oscillatory Ca2+ traces at 9mM glucose for wild type islets assessed by live-cell confocal microscopy. H) Three representative oscillatory Ca2+ traces at 9mM glucose for MOPR KO islets assessed by live-cell confocal microscopy. I) Oscillation frequency (oscillations/time) of individual Wild type and MOPR KO islets. MOPR KO islets exhibit significantly increased Ca2+ oscillation frequency ****P<0.0001 assessed by Mann-Whitney test n=101 wildtype islets n=106 MOPR KO islets from n=10 mice. J) Histogram showing the distribution of oscillation frequency of both wild type (gray) and MOPR KO (blue) islets. MOPR KO islets have an increased propensity to have high oscillation frequency.

Another key physiological measure of islet Ca^2+^ dynamics is Ca^2+^ oscillations ^29,37^. These oscillations are often reported to be proportional to insulin secretion ^30,38^, and deviations in oscillatory activity are known to have profound downstream effects on glucose metabolism ^39^. To assess differences in Ca^2+^ oscillations between MOPR KO and wild type islets, we incubated islets in 9mM glucose for an hour before commencing live cell Ca^2+^ imaging. MOPR KO islets often had more complex, compound-like oscillatory responses which have been previously reported ^29,35,37,40^. For reference, we included three representative traces each for wild type (Figure 3G) and MOPR KO islets (Figure 3H). When comparing all oscillating islets between wild type and MOPR KO genotypes, we found that MOPR KO islets had significantly increased oscillation frequency compared to wild type islets (Figure 3I). This increase was reflected as a noticeable qualitative modal shift in oscillation frequency among the population of islets (Figure 3J).

### Selective antagonism of the Mu opioid receptor increases Ca^2+^ accumulation but reduces oscillation frequency

Though effective, genetic deletion approaches can leave room for compensatory responses that are not reflective of true physiological function. As an alternative to the MOPR KO, we used the MOPR selective antagonist CTAP to dynamically block MOPRs on wild type islets during Ca^2+^ imaging experiments. To test the effects of CTAP in islet Ca^2+^ dynamics, we first imaged wild type islets at 3mM glucose. Next, we simultaneously added glucose to increase the glucose islet environment to 9mM in addition to either 10μM CTAP or vehicle. To assess relative uniformity in islets between the separate groups subsequently treated with either 10μM CTAP or vehicle, basal Ca^2+^ signal at 3mM glucose was compared. Basal signal between wild type islets treated with vehicle or CTAP was not different during the 3mM glucose baseline period (Figure 4A). Like genetic deletion of MOPR, following islet transition to 9mM glucose we observed an increased Ca^2+^ accumulation associated with CTAP treatment, which resulted in an increased area under the curve (Figure 4B-C). Analogous to MOPR KO islets, wild type islets treated with CTAP simultaneously with glucose addition significantly increased the elevation of Ca^2+^ signal when islets transitioned from 3mM glucose to 9mM glucose (Figure 4D).

**Figure 4.**
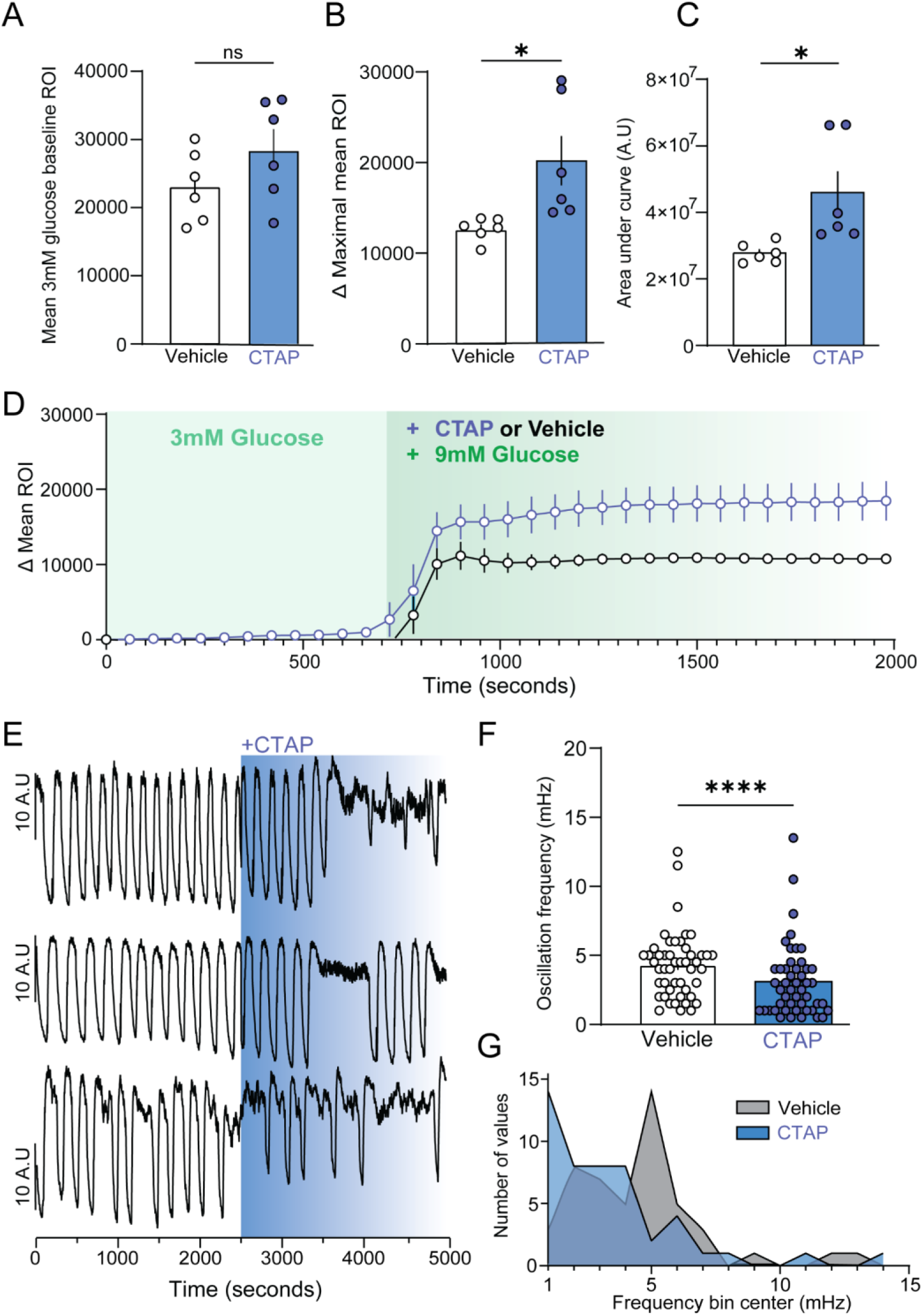
Mu opioid receptor antagonist CTAP increases islet Ca2+ accumulation when transitioning from 3mM glucose to 9mM glucose and decreases Ca2+ oscillation frequency. A) Baseline raw ROI signal at 3mM glucose of wild type islets before drug treatment. There is no significant difference between 3mM glucose baselines, n=6 mice for each condition, assessed by paired t-test P>0.05, error bars show +/- SEM. B) Maximal delta signal of Ca2+ accumulation in high glucose of wild type islets treated with 10μM CTAP or vehicle derived from D. CTAP significantly increases Ca2+ accumulation assessed by paired t-test *P<0.05 n=6, error bars show +/- SEM. C) Area under curve analysis from D of wild type islets treated with either CTAP or equivalent vehicle. CTAP significantly increases area under the curve assessed by unpaired t-test *P<0.05 n=6 error bars show +/- SEM. D) Ca2+ accumulation time course starting at 3mM glucose transitioned to 9mM at 750 seconds comparing wild type ex vivo islets treated with either 10μM CTAP or equivalent vehicle on board with glucose step. Images were quantified through mean region of interest (ROI) analysis of 20-25 islets per mouse from n=6 mice with Delta ROI normalized to signal from 3mM glucose at equilibrium. Error bars show +/- SEM. E) Representative Ca2+ oscillation trace of wild type islets at 9mM glucose subjected to vehicle (non-blue shaded) then 10μM CTAP at 2500 seconds (blue shaded). F) Oscillation frequency (oscillations/time) of individual islets treated with either vehicle or CTAP. Wild type islets exhibit decreased oscillation frequency following treatment of CTAP ****P<0.001 assessed by Willcoxon paired rank test n=48 islets from n=6 mice. G) Histogram showing the frequency distribution of oscillation frequency of wild type islets subjected to vehicle or CTAP. The modal Ca2+ oscillation associated with CTAP treatment is shifted left-ward.

To test CTAPs effect on Ca^2+^ oscillations, we incubated a separate set of wild type islets in 9mM glucose and recorded their Ca^2+^ oscillations for 2500 seconds to establish a baseline before treatment with 10μM CTAP. After CTAP treatment islets were imaged for an additional 2500 seconds. Surprisingly, unlike genetic deletion of MOPR, CTAP treatment decreased oscillation frequency (Figure 4E-F). This was reflected as modal shift across the population of treated islets, where the modal oscillation frequency associated with the CTAP treated group was decreased compared to vehicle control (Figure 4G).

### Selective agonism of Mu opioid receptor decreases Ca^2+^ accumulation but increases oscillation frequency

With prior observations on islet Ca^2+^ signaling with genetic deletion of MOPR and selective antagonism through CTAP treatment both increasing Ca^2+^ accumulation but having oppositional effects on Ca^2+^ oscillations, we sought to assess the effects of MOPR agonism on Ca^2+^ dynamics. Here we replicate the CTAP series of experiments in figure 4A-G but substituting 10μM CTAP for 10μM of MOPR full agonist DAMGO. As previously observed with CTAP, baseline signal at 3mM glucose prior to glucose addition with simultaneous treatment of either drug or vehicle was not significantly different (Figure 5A). When islets were transitioned to the 9mM glucose environment and treated with DAMGO, maximal Ca^2+^ accumulation was significantly decreased relative to vehicle and resulted in a decreased area under the curve (Figure 5B-C). The decreased Ca^2+^ accumulation associated with DAMGO treatment appeared to be sustained across the time course of the experiment (Figure 5D).

**Figure 5.**
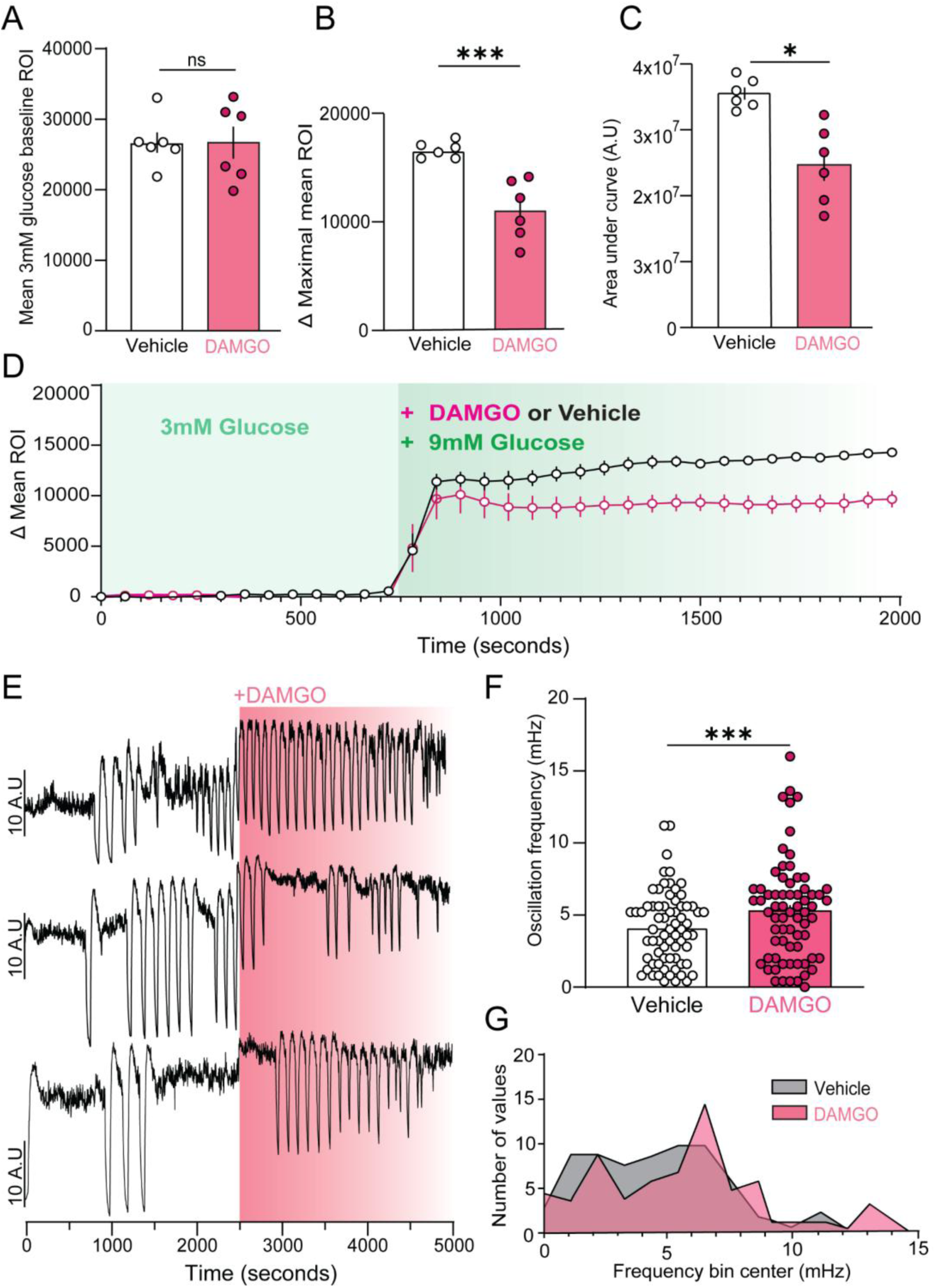
Mu opioid receptor agonist DAMGO decreases islet Ca2+ accumulation when transitioning from 3mM glucose to 9mM glucose and increases Ca2+ oscillation frequency at 9mM glucose. A) Baseline raw ROI signal at 3mM glucose of wild type islets before drug treatment. There is no significant difference between 3mM glucose baselines, n=6 mice for each condition, assessed by paired t-test P>0.05, error bars show +/- SEM. B) Maximal delta signal of Ca2+ accumulation in high glucose of wild type islets treated with 10μM DAMGO or vehicle derived from A. DAMGO significantly reduces Ca2+ accumulation assessed by paired t-test ***P<0.001 n=6, error bars show +/- SEM. C) Area under curve analysis of wild type islets treated with either DAMGO or equivalent vehicle. DAMGO significantly decreases area under the curve assessed by unpaired t-test *P<0.05 n=6 error bars show +/- SEM. D) Ca2+ accumulation time course starting at 3mM glucose transitioned to 9mM at 750 seconds comparing wild type ex vivo islets treated with either 10μM DAMGO or equivalent vehicle on board with glucose step. Images were quantified through mean region of interest (ROI) analysis of 20-25 islets per mouse from n=6 mice with Delta ROI normalized to signal from 3mM glucose at equilibrium. Error bars show +/- SEM. E) Representative Ca2+ oscillation trace of wild type islets at 9mM glucose subjected to vehicle (non-pink shaded) then 10μM DAMGO at 2500 seconds (pink shaded). F) Oscillation frequency (oscillations/time) of individual islets treated with either vehicle or DAMGO. Wild type islets exhibit increased oscillation frequency following treatment of DAMGO ***P<0.001 assessed by Willcoxon paired rank test n=69 islets from n=8 mice. G) Histogram showing the distribution of oscillation frequency of wild type islets subjected to vehicle or DAMGO. The modal Ca2+ oscillation associated with DAMGO treatment is shifted right-ward.

Given the differences between CTAP and vehicle and between MOPR KO and wild type islets on Ca^2+^ oscillations, we were intrigued to assess how MOPR agonism impacts islet Ca^2+^ oscillations. Here we adopt the same experimental setup we implemented to assess the effect of CTAP on Ca^2+^ oscillations in figures 4E-G but again substituted 10μM CTAP for 10μM DAMGO. Unlike CTAP, we observed an increase in Ca^2+^ oscillations associated with DAMGO treatment (Figure 5E-F). Although Ca^2+^ oscillation frequency increased in response to DAMGO treatment, towards the end of the trialing period, DAMGO appeared to disrupt oscillatory behavior, indicating a degeneration of regular oscillations (Figure 5E). When considering the whole population of islets, the occurrence of higher oscillation frequencies in the DAMGO treated islet population markedly shifted towards a greater modal frequency (Figure 5G).

### Mu opioid receptor antagonism decreases insulin secretion at 9mM glucose, but neither constitutive knockout of MOPR nor MOPR agonism affects insulin secretion

We next investigated glucose stimulated insulin secretion (GSIS) at 3mM glucose and 9mM glucose to assess whether constitutive knockout of MOPR or pharmacological modulation through CTAP and DAMGO treatments impacted static insulin secretion. The assay workflow is shown as a schematic (Figure 6A). Surprisingly, increased Ca^2+^ dynamics associated with genetic deletion of MOPR did not translate to increased insulin secretion and there was no significant effect on insulin secretion in both low- and high-glucose conditions compared to wild type controls (Figure 6B). Conversely, in line with Ca^2+^ oscillation data, 10μm CTAP treatment significantly decreased insulin secretion under high-glucose conditions. The effect of CTAP appeared to only be present at a higher concentration given that there was no significant difference between vehicle treatment and 1μm CTAP under both glucose concentrations. Despite DAMGO decreasing overall Ca^2+^ accumulation in response to glucose and increasing Ca^2+^ oscillations in previous experiments, neither 10μm or 1μm DAMGO had any significant effect on insulin secretion at either 3mM or 9mM glucose. Considering the relationship between islet Ca^2+^ signaling (particularly Ca^2+^ oscillation frequency) and insulin secretion, there appears to be consistency with 10μm CTAP treatment causing both decreased Ca^2+^ oscillation frequency and decreased insulin secretion at 9mM glucose. However, genetic deletion of MOPR and DAMGO impacted Ca^2+^ dynamics but neither translated into any significant change in insulin secretion.

**Figure 6.**
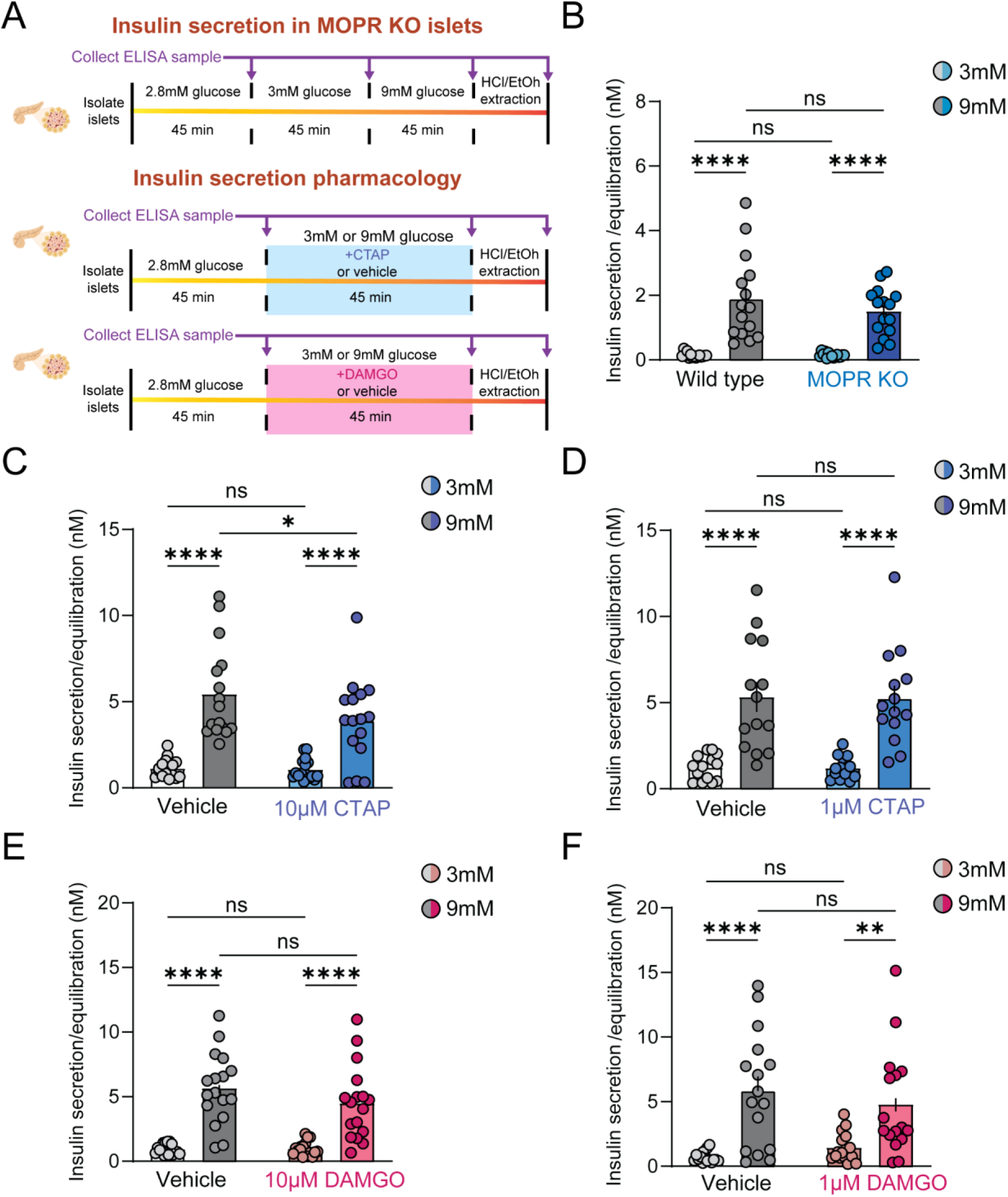
10μM CTAP significantly decreases insulin secretion under high glucose conditions, but constitutive knockout of mu opioid receptor and DAMGO has no effect on static insulin secretion. A) Schematic and workflow of static insulin secretion assay. Islets are equilibrated at 2.8mM glucose before drug treatments and transitioning to the final glucose concentration with ELISA samples extracted at each change of condition. All values reflect normalized values relative to total insulin content B) Insulin secretion of wild type (gray) or MOPR KO (blue) islets under either 3mM or 9mM glucose conditions. Constitutive knockout of MOPR has no significant effect on insulin secretion, assessed by 2-way ANOVA with Fisher’s least significant difference correction for multiple comparisons, n=15 mice P>0.05. C) Insulin secretion of wild type islets treated with either vehicle (gray) or 10μM CTAP at both 3mM and 9mM glucose. CTAP significantly decreases insulin secretion at 9mM glucose compared with vehicle assessed by 2-way ANOVA with Fisher’s least significant difference correction for multiple comparisons, n=16 mice *P<0.05. D) Insulin secretion of wild type islets treated with either vehicle (gray) or 1μM CTAP at both 3mM and 9mM glucose. 1μM CTAP has no significant effect on insulin secretion compared with vehicle assessed by 2-way ANOVA with Fisher’s least significant difference correction for multiple comparisons, n=14 mice *P>0.05. E) Insulin secretion of wild type islets treated with either vehicle (gray) or 10μM DAMGO (pink) at both 3mM and 9mM glucose. DAMGO has no significant effect on insulin secretion assessed by 2-way ANOVA with Fisher’s least significant difference correction for multiple comparisons, n=17 mice, P>0.05. F) Insulin secretion of wild type islets treated with either vehicle (gray) or 1μM DAMGO (pink) at both 3mM and 9mM glucose. 1μM DAMGO has no significant effect on insulin secretion assessed by 2-way ANOVA with Fisher’s least significant difference correction for multiple comparisons, n=17 mice, P>0.05.

### Mu opioid receptor knockout increases glucagon secretion at 3mM glucose, while CTAP and DAMGO have no effect

Lastly, similarly to glucose-stimulated insulin secretion, we investigated static glucagon secretion at both 3mM glucose and 9mM glucose conditions to assess whether constitutive knockout of MOPR or pharmacological modulation impacted static glucagon secretion. As with figure 6A we provide a schematic for the workflow of this assay (Figure 7A). Genetic deletion of MOPR appeared to significantly increase glucagon secretion at 3mM glucose (Figure 7B). Critically, the enhancement of glucagon secretion associated with MOPR KO islets was restricted to 3mM glucose conditions, where glucagon secretion would be physiologically stimulated. In contrast to glucose-stimulated insulin secretion data in figure 6C, CTAP treatment had no significant effect on glucagon secretion at every concentration trialed (Figure 7B). Similarly to insulin secretion experiments, DAMGO also appeared to have no significant effect on glucagon secretion at any of the concentrations trialed.

**Figure 7.**
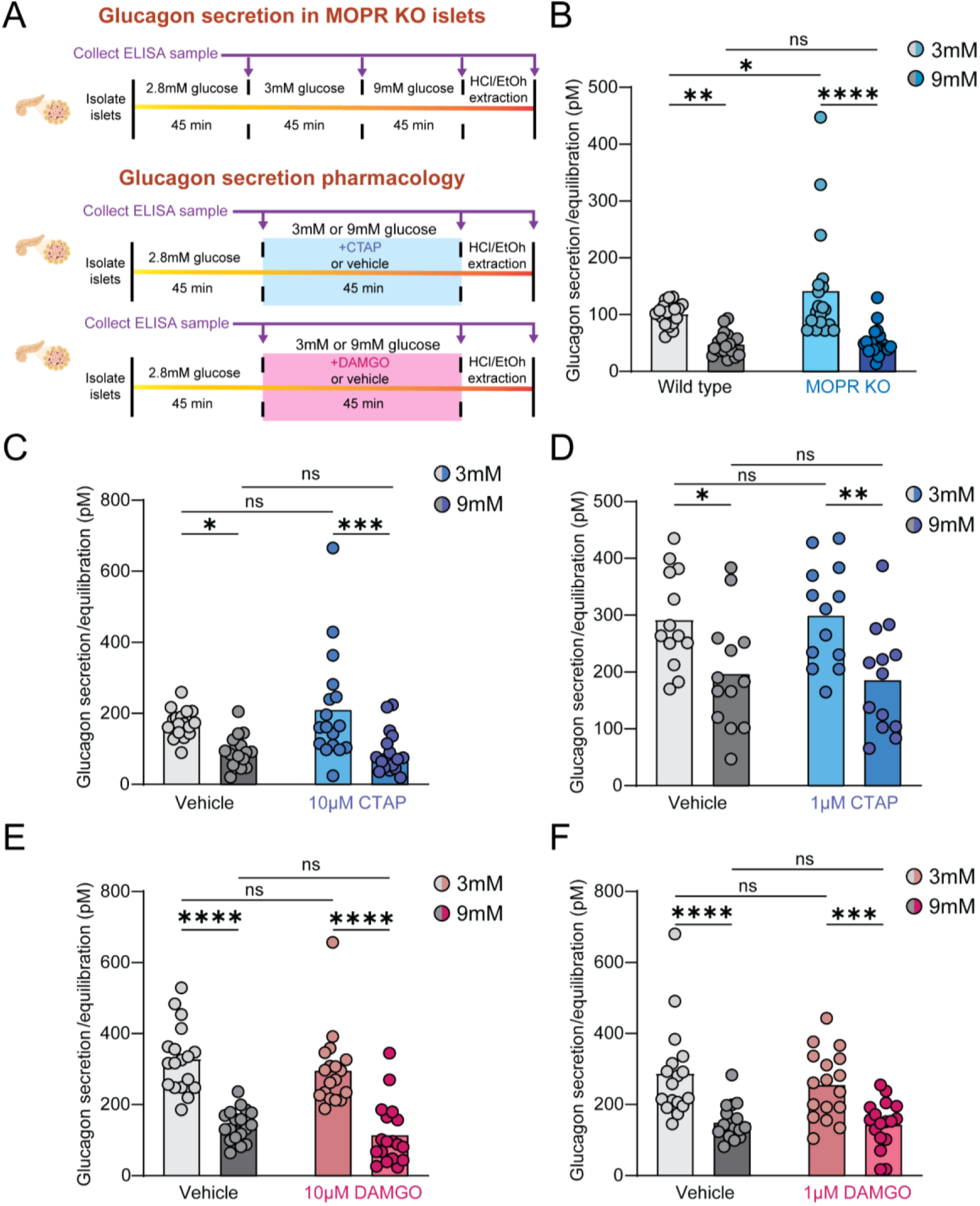
Constitutive mu opioid receptor knockout increases glucagon secretion under low glucose conditions, CTAP and DAMGO have no effect on glucagon secretion. A) Schematic and workflow of static glucagon secretion assay. Islets are equilibrated at 2.8mM glucose before drug treatments and transition to the final glucose concentration with ELISA samples extracted at each change of condition. All values reflect normalized values relative to total glucagon content. B) Glucagon secretion of wild type (gray) or MOPR KO (blue) islets under either 3mM or 9mM glucose conditions. Constitutive knockout of MOPR significantly increased glucagon secretion at 3mM glucose, assessed by 2-way ANOVA with Fisher’s least significant difference correction for multiple comparisons, n=19 mice *P<0.05. C) Glucagon secretion of wild type islets treated with either vehicle (gray) or 10μM CTAP at both 3mM and 9mM glucose. CTAP has no significant effect on glucagon secretion compared with vehicle, assessed by 2-way ANOVA with Fisher’s least significant difference correction for multiple comparisons, n=17 mice P>0.05. D) Glucagon secretion of wild type islets treated with either vehicle (gray) or 1μM CTAP at both 3mM and 9mM glucose. 1μM CTAP has no significant effect on insulin secretion compared with vehicle assessed by 2-way ANOVA with Fisher’s least significant difference correction for multiple comparisons, n=13 mice P>0.05. E) Glucagon secretion of wild type islets treated with either vehicle (gray) or 10μM DAMGO (pink) at both 3mM and 9mM glucose. DAMGO has no significant effect on insulin secretion assessed by 2-way ANOVA with Fisher’s least significant difference correction for multiple comparisons, n=18 mice, P>0.05. F) Insulin secretion of wild type islets treated with either vehicle (gray) or 1μM DAMGO (pink) at both 3mM and 9mM glucose. 1μM DAMGO has no significant effect on insulin secretion assessed by 2-way ANOVA with Fisher’s least significant difference correction for multiple comparisons, n=17 mice, P>0.05.

## Discussion

Expression of endogenous opioids and their respective receptors in the endocrine pancreas has been met with skepticism but have been validated with a variety of approaches mainly focused on single cell transcriptomics and RNA approaches. To date, there is evidence for the MOPR and delta opioid receptor expression in the pancreatic islet ^21,41,42^. More recently, relative expression of the *Oprm1* gene in both mouse and human islets has been established using qPCR and fluorescent in situ hybridization and has demonstrated robust evidence for pancreatic MOPR expression. In addition, MOPR expression appears dysregulated in disease states as observed by decreased MOPR transcription in response to type 2 diabetes in humans and obesity mice ^21^. To support existing evidence for pancreatic MOPR expression, we used three different approaches to rigorously validate expression. First, we analyzed single-cell RNA sequencing to show *Oprm1*, and its endogenous agonist precursors (pro-enkephalin and pro-opiomelanocortin), were actively transcribed in islet cells, albeit with low detection for *Oprm1* and *Penk* in this approach. Second, we used in situ hybridization to assess the architecture of *Oprm1* transcription in whole islet samples and through co-localization analysis with specific islet cell type markers. We determined that *Oprm1* is expressed in approximately 50% of islet cells with no obvious expression proclivity for any islet cell type. Finally, while our data and previous publications indicate *Oprm1* RNA is expressed in islets ^19^, evidence for protein expression has remained elusive. Using immunoblotting with a well validated MOPR specific antibody, we show that MOPR protein is expressed in pancreatic primary islets ^43–45^.

Following our expression studies, we next determined the functional capabilities of MOPRs in whole islets. To interrogate Gαi/o canonical MOPR signaling, we used a FRET based approach to measure changes in cAMP accumulation. We found that DAMGO suppressed cAMP accumulation, and was reversible through pretreatment with CTAP, indicating pharmacologically responsive MOPRs in mouse islets. Strikingly, MOPR also appears to have some influence on how islets respond to glucose. Specifically, wild type islets appeared to have an inversely proportional relationship between glucose concentration and cAMP accumulation, which was then disrupted by MOPR KO in high glucose conditions. has been observed in dynamic TIRF microscopy. Although similar cAMP/glucose concentration relationships have been previously reported, this relationship does not appear consistent and varies dependent on the assay utilized to measure cAMP accumulation in whole islets ^46–49^. For example, our method used an end point assay to measures signal from the whole islet lysate, leaving unresolved whether there may be dynamic fluctuations in cAMP across time or cell types in addition to concentration. Future work using live-cell FRET imaging with either a cAMP sensor directly, or a cAMP dependent sensor downstream, such as a protein kinase A, would have the added benefit of recording dynamic cAMP signaling.

To investigate how MOPRs regulate dynamic Gβγ signaling in response to glucose, we used live-cell confocal microscopy with the Ca^2+^ sensitive dye Calbryte^TM^ 520 AM. We showed that genetic deletion of MOPR increases Ca^2+^ accumulation, both under low and high glucose conditions (Figure 3C-F). MOPR KO islets also exhibited increased oscillation frequency at 9mM glucose compared with wild type islets. Pharmacological antagonism of MOPR via CTAP similarly increased overall Ca^2+^, but surprisingly decreased oscillation frequency. This suggests that there could be long term plastic changes in cellular physiology or signaling in KO islets. One contributor to these long-term changes could be cAMP. Since MOPR KO islets exhibit an increased cAMP accumulation phenotype at high glucose conditions, this may cause changes to protein expression through cAMP response element binding proteins (CREB) that are responsible for this compensation mechanism^50,51^. In islets, increases in cAMP accumulation has been shown to have long term cellular effects such as increasing β cell proliferation while suppressing apoptosis and increasing insulin synthesis ^52^. Regardless of the precise mechanistic differences between genetic and antagonist results, the impact of MOPR disruption of Ca^2+^ dynamics indicates a potential endogenous paracrine mechanism wherein islet expressed MOPRs are physiologically activated by endogenously expressed agonists. Our data and other sequencing data support this hypothesis, as both enkephalin and proopiomelanocortin appear to be expressed in islets. Further studies focusing on these peptides would be of great interest moving forward.

Like CTAP, MOPR agonism via DAMGO had paradoxical effects on Ca^2+^. Although overall Ca^2+^ accumulation was suppressed (consistent with downregulation of Gβγ signaling after Gαi/o activation), Ca^2+^ oscillations were increased. However, it should be appreciated that while DAMGO increased oscillations overall, the amplitude of these oscillations appeared to degrade over time. The degeneration of Ca^2+^ oscillations despite transiently increasing frequency may partly explain how Gi/o GPCR activation reduces insulin secretion given the positive correlation between Ca^2+^ oscillations and insulin secretion. Given the complex signaling effects of opioid modulation and islet physiology, future research investigating clinically utilized ligands that target MOPR will be critical for determining how their specific administration may influence metabolic and glycemic activity.

To compliment the islet signaling data we have presented, we also assessed how constitutive knockout of MOPR and modulation of MOPR pharmacologically impacted hormone secretion. In line with reduced Ca^2+^ oscillation frequency, treatment with 10μM CTAP decreased insulin secretion compared with vehicle-treated islets at high glucose. Genetic deletion of MOPR however had no effect on insulin secretion despite increasing both Ca^2+^ accumulation and Ca^2+^ oscillation frequency. These non-parallel observations suggest that not only does MOPR integrate with canonical intracellular signaling cascades associated with hormone secretion but measurably tunes insulin secretion when acutely, but not chronically, attenuated. The impact of genetic deletion and antagonism of MOPR was mirrored in glucagon secretion experiments where treatment with CTAP did not affect glucagon secretion while genetic deletion of MOPR increased glucagon secretion under low glucose conditions. When glucagon and insulin secretion data are considered together, it appears that acute attenuation of MOPR impacts beta cell activity whereas chronic MOPR attenuation through constitutive knockout impacts alpha cell activity. Despite glucagon secretion enhancement by constitutive MOPR KO under low glucose conditions, MOPR modulation appears to exhibit most of its influence on intracellular signaling when the islets are under high glucose concentrations. This was also the case in previous studies investigating glucagon secretion ^19,21^ and in figure 2, where cAMP accumulation was only increased in MOPR KO islets when the islets were incubated at either 6mM or 9mM glucose. Considering previous studies, future work should further investigate how MOPR function may be uniquely recruited on different cell types to impact islet activity. However, it should be appreciated that in most clinical conditions (opioid agonists for pain relief, antagonists for overdose treatment), these non-specific effects are likely more informative.

Given that in this study and others ^53^ found evidence for endogenous opioid peptide expression (including beta endorphin, enkephalin, and dynorphin), mechanisms of their potential secretion and subsequent paracrine signaling, along with islet cell type specific effects should be explored. Similarly, it would be intriguing to investigate whether endogenous opioid peptide agonists are secreted under high glucose conditions which under physiological conditions would activate pancreatic MOPRs but are restricted under both conditions of selective antagonism and genetic deletion of MOPR. However, since the outcomes of MOPR selective antagonism and genetic deletion of MOPR are different with respect to intracellular Ca^2+^ dynamics and secretion, this points to a complex relationship between intact islet physiology and the integration of islet cell type specific MOPR signaling with islet second messenger effectors tied to hormone secretion.

In sum, these experiments demonstrate a complex, yet generally consistent effect of MOPRs on islet physiology and secretion. It is likely that some of the observed complexity/paradoxes are driven by concurrent modulation of MOPRs on different and antagonistic cell types (alpha versus beta, delta versus beta). The approaches used in this study were unable to delineate cell type specific effects, thus future work will necessarily require cell type selective targeting to better assess the role of MOPRs on each cell type. However, the lack of cellular specificity is likely more translational, as exogenously delivered opioids like fentanyl, oxycodone, and tramadol will act on all available MOPRs. Since exogenous opioids that target the MOPR span a wide spectrum of efficacy including antagonists, partial, full and biased agonists ^54–56^, opioid use in different circumstances may have both different, and inadvertent consequences on metabolism and glucose homeostasis. Therefore, a balanced approach will be required to understand both the molecular and functional effects of opioids across a wide domain of contexts.

## Acknowledgements and author contributions

Michael Keith designed/performed experiments and wrote the manuscript. Christine Stander and Diego De Gregorio researched data. Andy Huang aided Michael Keith in researching data. Sepideh Sheybani-Deloui researched data, Shannon Townsend, Jeffrey M Zigman and Jing W Hughes advised on experiments and reviewed/edited the manuscript. Daniel C Castro supervised experimental work, interpreted data and reviewed/edited the manuscript.

---

**Figure S1.**
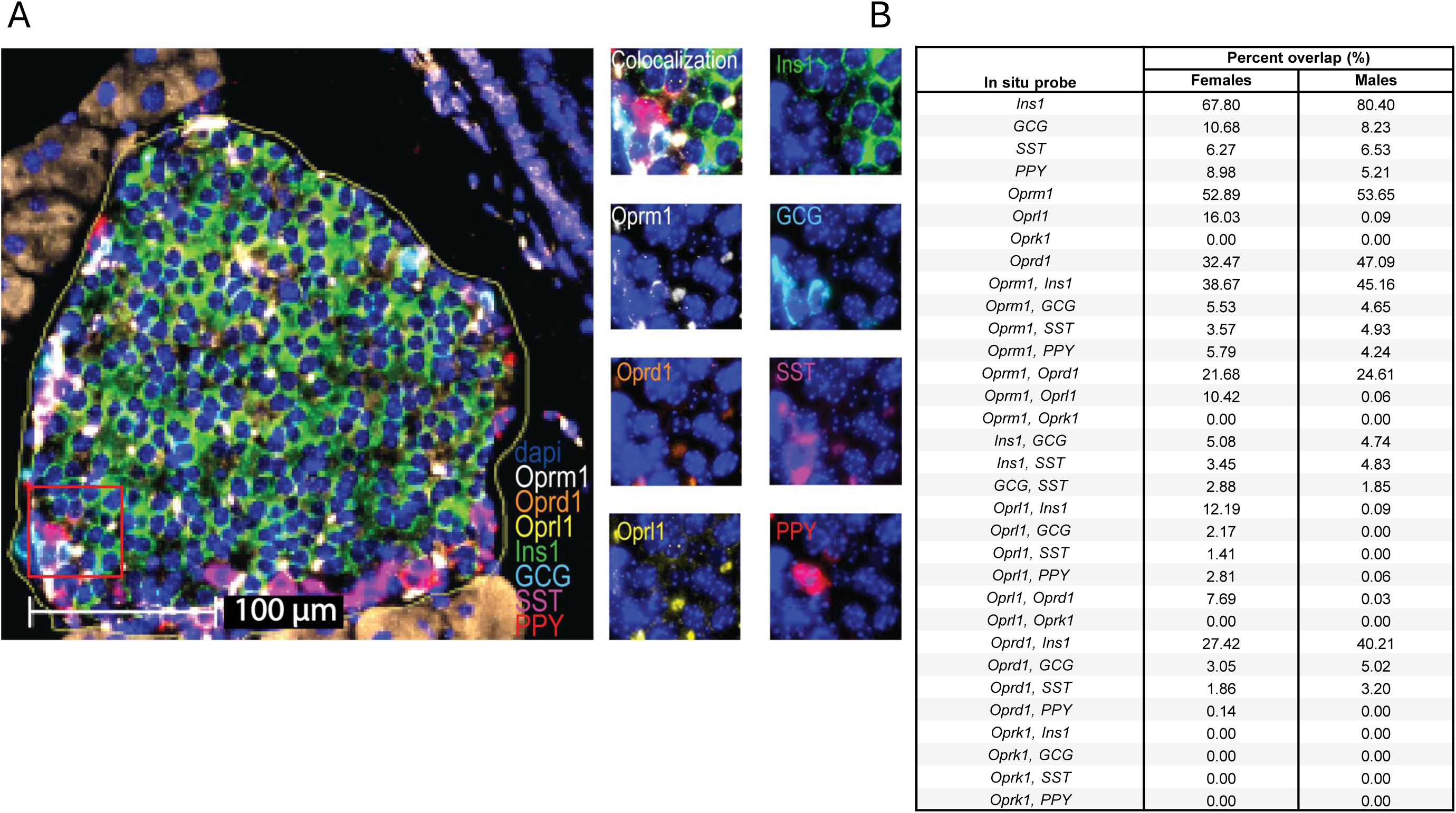
Delta and nociceptin opioid receptors are transcribed in the murine islet of Langerhans. **A)** Isolated wild type islet RNA scope image showing individual and merged transcripts of opioid receptors including mu opioid receptor (OPRM1, white), delta opioid receptor (Oprd1, orange), nociceptin receptor (Oprl1, yellow) in addition to insulin, (Ins1, green) glucagon, (Gcg, blue) somatostatin, (Sst, pink) and polypeptide Y (PPY, red) **Scale = 100μm** **B)** Table showing percent overlap of opioid receptor expression (Orpm1, Oprd1, Oprk1, Oprl1) with pancreatic peptides (Ins1, GCG, SST, PPY) in islets from mice (n=3/sex). Values represent a mean percent overlap of marker co-expression.

**Figure S2.**
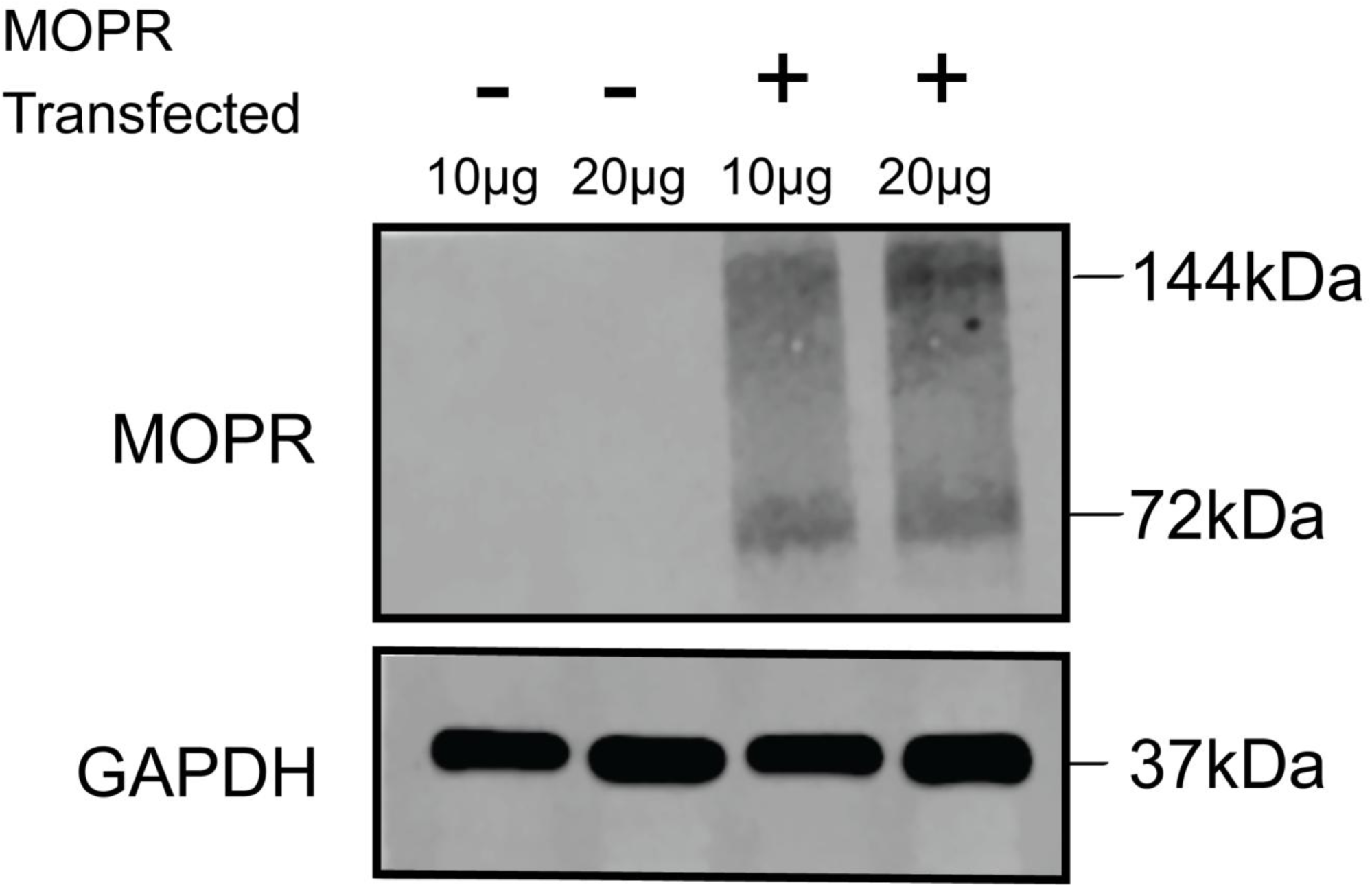
HEK293-MOPR transient expression system validates antibody specificity for use in ex-vivo islets. Representative western blot of HEK293 cell lysates transiently transfected with either the mu opioid receptor (+) or equivalent empty vector (-) immunoblotted for the Mu opioid receptor (upper panel) and GAPDH for loading control (lower panel). Blots were loaded with either 10μg or 20μg total protein, determined with BCA protein assay.

**Figure S3.**
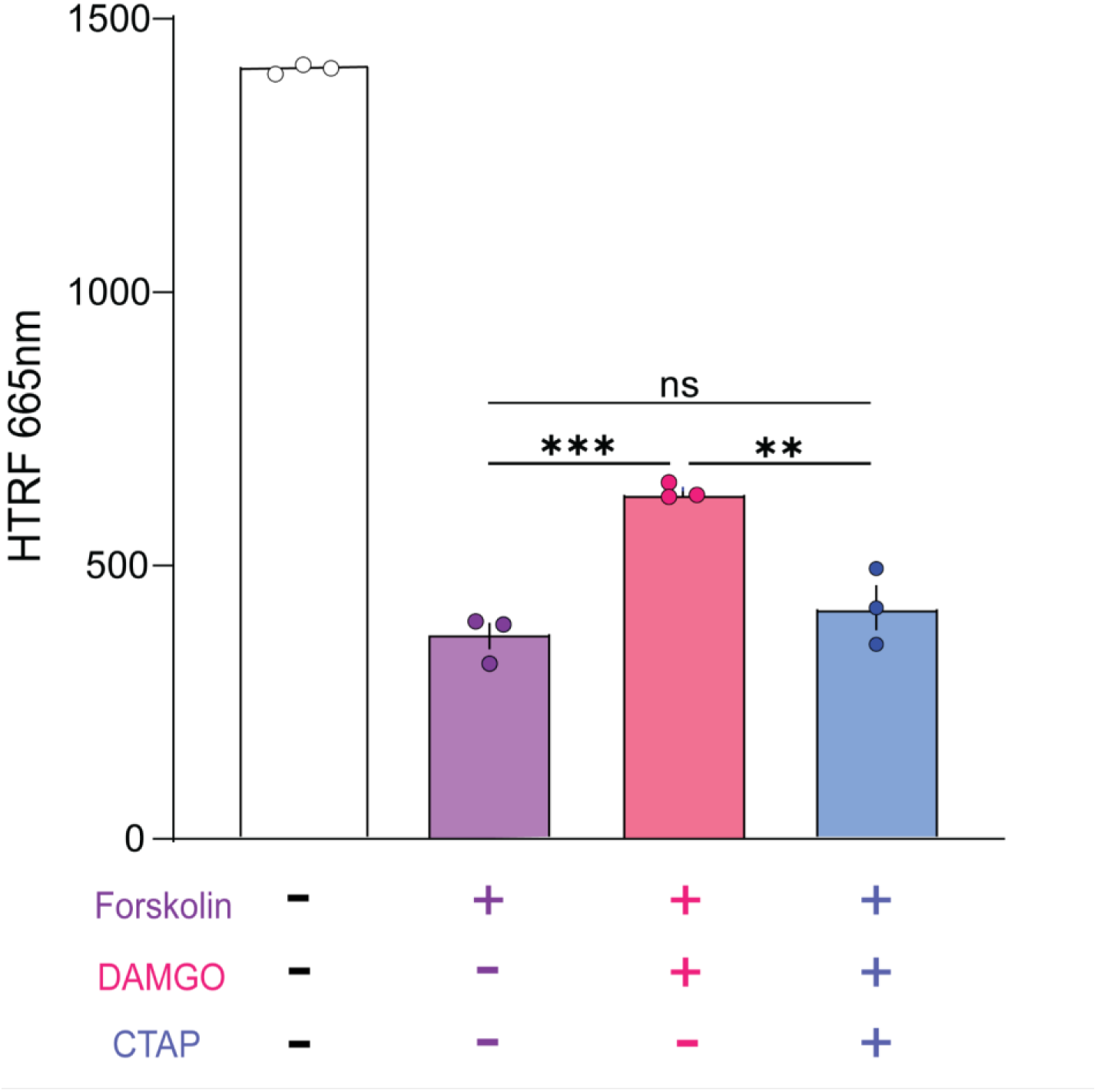
MOPR cAMP pharmacology in HEK293 expression system. HEK293 cells transiently transfected with the mu opioid receptor respond analogously to ex-vivo isolated islets when treated with MOPR ligands. HEK293-MOPR cells (2000 cells per well) subjected to either vehicle control (white), 10μM forskolin (purple), 10μM forskolin and 10μM DAMGO (pink) or 10μM forskolin, 10μM DAMGO and 10μM CTAP (blue). DAMGO significantly suppressed forskolin induced cAMP accumulation, forskolin (purple) vs forskolin + DAMGO (pink) ***P<0.001. CTAP blocks the effect of DAMGO suppressing forskolin induced cAMP accumulation, forskolin + DAMGO (pink) vs forskolin + DAMGO + CTAP (blue) **P<0.01. Due to antagonism through CTAP, there is no significant difference in cAMP accumulation between forskolin alone (purple) vs forskolin + DAMGO + CTAP (blue) P>0.05. Statistical differences were assessed by one way ANOVA with Bonferroni’s test for multiple comparisons, each data point shows the mean from n=3 independent experiments, with error bars showing +/- SEM.

