## Supplemental materials for "MOPRs in mouse islets of Langerhans modulate cell signaling and secretion"

Supplemental methods

Human embryonic kidney (HEK293T) cell culture and transfection

HEK293T cells were cultured in 100mm cell culture dishes (Corning, 353003) with 10mL Dulbecco's modified essential media supplemented with 10% fetal bovine serum and 1% penicillin/streptomycin (HEK293T cell culture media) and incubated at 37ºC, 5% CO2 in a humidified incubator. For transient transfection to express the MOPR, an 80% confluent dish was incubated with 2μg of plasma encoding receptor construct in pcDNA3.1 with lipofectamine according to manufacturer's protocol. In short, cells were incubated with DNA/lipofectamine 2000 complexes for 6 hours before replacement with fresh HEK293T cell culture media. Following an additional 48-hour incubation, transfected HEK293T cells were utilized in the cAMP assay as a positive control for MOPR activity.

Preparation of HEK293-MOPR cell lysates for western blot (supplemental)

HEK293 cells were transiently transfected with MOPR plasmid or equivalent empty vector as described above. Following transfection, cells were washed with ice cold PBS twice and collected through mechanical disruption using a cell scrapper (Corning, 29442-202) and lysed using 1mL of ice-cold RIPA buffer supplemented with protease cocktail inhibitor, 0.1mM PMSF and incubated for 30 minutes on ice, as previously described above. Finally, HEK293-MOPR cell lysates protein content was determined using BCA protein assay, separated using SDS-PAGE, transferred to PVDF, blocked and immunoblotted for MOPR in addition to GAPDH as loading control using the same methodology as ex-vivo islets as described above.

cAMP accumulation assay in HEK293 cells transiently transfected with MOPR

For cAMP accumulation assays conducted in HEK293 cells, transfections were performed as described above and disassociated at 37ºC, 5% $CO_2$, using 1mL of 0.5mM EDTA (Sigma, E9884). Cells were washed by resuspending in PBS and centrifuging at 300g for 5 minutes. To determine the concentration of cells, the cell pellet was dissolved in 5mL PBS and 10μL of cell suspension and 10μL trypan blue dye was dispensed in Countess™ cell counting chamber slides. Three slides were prepared and loaded into Countess automated cell counter to determine cellular concentration. After determining the cellular concentration through taking the mean of the cell counter readout, the cell solution was diluted to 400 cells/μL using stimulation buffer to dispense 2000 cells per well via 5μL additions. Dispensed HEK293-MOPR cells were treated with vehicle, forskolin, DAMGO and CTAP analogously to ex vivo isolated islets before incubating for 1 hour at room temperature with cAMP detection reagents as previously described.
